# Cryptogams signify key transition of bacteria and fungi in Arctic sand dune succession

**DOI:** 10.1101/699876

**Authors:** Heli Juottonen, Minna Männistö, Marja Tiirola, Minna-Maarit Kytöviita

## Abstract

- Primary succession models focus on aboveground vascular plants. However, the prevalence of mosses and lichens, i.e. cryptogams, suggests they play a role in soil successions. Here, we explore whether effects of cryptogams on belowground microbes can facilitate progressive shifts in sand dune succession.
- We linked aboveground vegetation, belowground bacterial and fungal community, and soil chemistry in six successional stages in Arctic inland sand dunes: bare sand, grass, moss, lichen, ericoid heath and mountain birch forest.
- Compared to the bare sand and grass stages, microbial biomass and the proportion of fungi increased in the moss stage, and later stage microbial groups appeared despite the absence of their host plants. The microbial communities of the lichen stage resembled the communities in the vascular plant stages. Bacterial community correlated better with soil chemistry than with vegetation, whereas the correlation of fungi with vegetation increased with vascular vegetation.
- Distinct bacterial and fungal patterns of biomass, richness, and plant-microbe interaction showed that the aboveground vegetation change structured the bacterial and fungal community differently. The nonalignment of aboveground vs. belowground changes suggests that cryptogams can drive succession towards vascular plant dominance through microbially mediated facilitation in eroded Arctic soil.

## Introduction

Development of vegetation on bare land and the sequence of vegetative stages that predictably follow each other have been described in several theories of succession (Egler, 1954; Wilson *et al*., 1992). According to widely accepted views, primary succession is influenced by site abiotic conditions (climate, soil parent material, topography) and biotic factors (plant species pool present, order of arrival, species interactions). Although focusing on plants, most models of terrestrial primary succession overlook bryophyte communities (e.g. Connell and Slatyer, 1977; Baker and Walford, 1995), whereas most reports of primary succession include moss and lichen dominated stages (Chapin *et al*., 1994; Lichter, 1998; Hodkinson *et al*., 2003; Jones and Henry, 2003).

Ecosystem succession includes stochastic elements for instance through priority effects and, on the other hand, deterministic influences such as competition and facilitation (Måren *et al*., 2018). Cryptogams, here mosses and lichens, may have diverse but unexplored competitive and facilitative effects on the development of vascular plant and belowground microbial communities. Firstly, dispersal of cryptogams by spores minimizes dispersal limitation – a feature classically included in models explaining succession of vascular plant communities (Makoto and Wilson, 2019). The rapid arrival of mosses warrants them a priority effect in establishing plant communities. Secondly, although mosses and lichens lack roots and root-mediated effects on microbial communities, they affect soil temperature, moisture, and carbon and nitrogen availability, which may promote or inhibit vascular vegetation (Van der Wal and Brooker, 2004; Gornall *et al*., 2007; Cornelissen *et al*., 2007; Gornall *et al*., 2011). Moss rhizoids and litter provide carbon to the belowground soil (Bowden, 1991), and mosses can release carbon also in drying and wetting cycles (Wilson and Coxon, 1999). Many lichens characteristic of primary succession fix atmospheric nitrogen in specific structures called cephalodia (Vitousek, 1994), whereas moss leaves are a habitat of N_2_-fixing bacteria (DeLuca *et al*., 2002; Arróniz-Crespo *et al*., 2014). Thirdly, mosses and lichens produce unique secondary metabolites that may have adverse effects on vascular plants and beneficial or adverse effects on microbes (Cornelissen *et al*., 2007; Xie and Lou, 2009).

Despite the fact that microbes are the first colonizers of any barren surface and modify soil chemistry during succession, there are only few models predicting successional trajectories of bacteria and fungi (Jackson, 2003; Fierer *et al*., 2010 Dini-Andreote *et al*., 2015; Tripathi *et al*., 2018; Ortiz-Álvarez *et al*., 2018). Belowground microbial community shifts in early succession are considered to be driven by soil chemistry, external resources and dispersal of microbes, and in later stages by biotic factors, such as establishment and changes in the vegetation (Brown and Jumpponen, 2014; Jiang *et al*., 2018). Bacterial versus fungal communities can be expected to show distinct trajectories in primary succession: Bacteria are smaller and believed to disperse easier than fungi, which may lead to more deterministic community assembly for bacteria (Schmidt *et al*., 2014; Powell *et al*., 2015) and higher influence of priority effects and stochastic effects for fungi (Brown and Jumpponen, 2014; Schmidt *et al*., 2014; Jiang *et al*., 2018). Further, the physiological diversity in bacteria enables some of them to survive as autotrophs in bare oligotrophic soils of early successional stages in the absence of plants fixing carbon (Nemergut *et al*., 2007; Schmidt *et al*., 2008; Duc *et al*., 2009). Fungi in these habitats depend on already fixed carbon and nitrogen from wind-blown, ancient, or microbially fixed sources (Schmidt *et al*., 2014). The critical shift to an ecosystem based on the biomass production by the resident plants is associated with changes in the microbial community (Bardgett and Walker, 2004; Edwards *et al*., 2006; Blaalid *et al*., 2012; Knelman *et al*., 2012). Along succession, the fungi to bacteria ratio generally increases as soil organic matter increases and pH decreases (Pennanen *et al*., 2001; Tscherko *et al*., 2004; Bardgett and Walker, 2004). These results do not, however, fully predict vegetation development in succession. We suggest that belowground changes associated with cryptogams are a key factor in the ecosystem shift from early to late succession. We link belowground community development with that aboveground and expect the interaction strength to vary and that the community changes do not take place in complete synchrony as is inferred in facilitation models (Brooker *et al*., 2008).

Ecosystems are declining due to soil erosion and loss of vegetation cover (Montgomery, 2007; Vanwalleghem *et al*., 2017), and it is critical to identify the key factors promoting shifts between non-vegetated and vegetated stages in succession. Identifying microbial and plant communities that signal for the shift from autotrophic to mainly heterotrophic microbial biomass production can help identify factors that drive the shift back to vascular plant cover, soil stabilization, and a productive ecosystem. One example of areas where vegetation has been lost due to environmental changes and lack of recovery are Arctic inland sand dunes. The dunes have formed from wind-blown sand after the last ice age (Koster, 1988). Plants colonized the dunes later on, but today large areas lack vascular plant cover. Forest fires, insect outbreaks, and overgrazing may have caused the ecosystem degradation (Seppälä, 1995). In our study site in Northern Finland, wind erosion has carved deflation basins into the dunes. Inside the basin, primary succession vegetation stages have developed, and dune slopes surrounding the basins are covered by mountain birch forest, the climax stage in the present climate. Eroding inland sand dunes provide an excellent opportunity to study primary succession due to the mosaic of closely situated successional stages within deflation basin, including moss- and lichen-dominated areas. In temperate inland dunes, the cryptogam stages have been identified as pioneer vegetation stages of the succession (Sparrius *et al*., 2012). Previous work comparing the start and end points of the Arctic inland dune succession proposed that bacteria and fungi may show distinct successional patterns (Poosakkannu *et al*., 2017). The role of the cryptogam stages as part of successional trajectories above- and belowground has not been elucidated, although the prevalence of lichens and mosses suggests they are important.

In this study, we explored whether the belowground and aboveground community changes take place in synchrony in Arctic inland sand dune succession. We hypothesized that, compared to bare soil, the cryptogam-covered stages contain belowground soil microbial community more similar to the stages covered by vascular plants. Such nonalignment of belowground vs. aboveground communities could indicate that the cryptogam stages drive succession towards vascular plant dominance through microbially mediated facilitation. Secondly, we compared the belowground responses of two microbial groups – bacteria and fungi - to successional stages defined by aboveground vegetation. We hypothesized that i) in comparison to fungi, bacteria would be more influenced by cryptogam-associated changes in the soil chemical environment because of their limited saprophytic capacity and potential autotrophy, and ii) as powerful and obligate decomposers, fungal community would respond particularly to the appearance of vascular vegetation and consequently organic matter accumulation. Such a pattern would indicate differences in the way vegetation change and plant-microbe interaction structure bacterial vs. fungal communities along the succession. By linking the aboveground vegetation, the belowground microbial communities, and the associated soil chemical environment, we aimed to define the multi-faceted role of cryptogams in Arctic ecosystem succession.

## Materials and methods

### Study site and sampling

The study site is located in an inland aeolian (i.e. wind-deposited) dune area in Northern Fennoscandia (68° 29′ 16″ N, 24° 42′ 13″ E). The 1981–2000 average annual temperature in the region was −1.3 °C, with extreme air temperatures between +30 °C and −52 °C and the average annual precipitation 550 mm (Pirinen *et al*., 2012). We defined six successional stages in the dunes based on vegetation (Fig. 1a). Although the soil in all the stages was aeolian sand, in three stages the largest part of the surface was mostly exposed sand (92-100% sand), while in the other three stages vegetation covered the sand (0-52 % sand). The sand-exposed stages (‘sandy stages’) were: 1) sand: bare loose sand without plants or visible soil crust, 2) grass: bare loose sand with the grass *Deschampsia flexuosa* and without visible soil crust, and 3) moss: bare loose sand with the moss *Polytrichum piliferum*. The stages with extensive vegetation cover (‘vegetated stages’) were 4) lichen: the main species lichens *Stereocaulon* spp. and *Cladonia* spp., and the mosses *P. piliferum* and *Racomitrium ericoides*, 5) heath: the dominant species dwarf shrub crowberry *Empetrum nigrum* with *Cladonia* spp. and/or *Stereocaulon* spp., and 6) forest: mountain birch (*Betula pubescens* subsp. czerepanovii) forest, the main species in study plots the shrubs *E. nigrum* and *Vaccinium vitis-idaea*, the moss *Pleurozium schreberi*, *D. flexuosa*, and the herb *Linnaea borealis*. The sand, grass, moss, and lichen stages are located at the bottom or on the inner slopes of the deflation basins and heath at the edge of the basins. The remnant mountain birch forests are located around the basins at the crest of the dune or on the outer slope. Permanent study plots of 1 × 1 m were established in 2008 and included each of the six vegetation stages in seven deflation basins. The vegetation in the plots was recorded in detail in 2009 by point frequency method (100 points within 50 × 50 cm, presence/absence and frequency of species).

**Figure 1.**
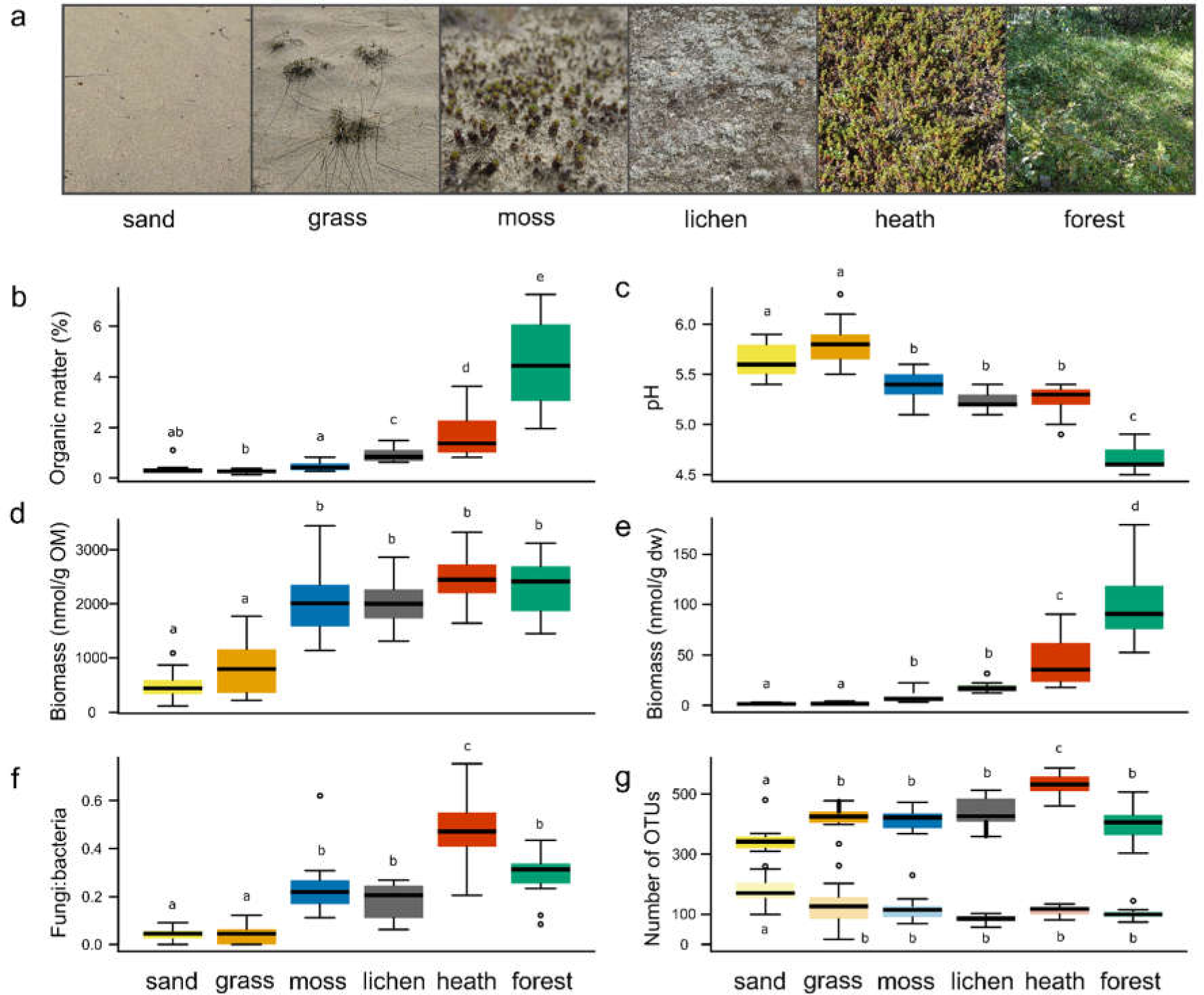
(a) Six successional stages defined in deflation basins of an Arctic inland sand dune area, (b) organic matter, (c) pH, (d) microbial biomass based on phospholipid fatty acids (PLFAs) per soil organic matter (OM) content, (e) microbial biomass per gram soil dry weight (dw), (f) ratio of fungal to bacterial biomass based on PLFA, and (g) bacterial 16S rRNA gene (upper row, darker colours) and fungal ITS amplicon (lower row, lighter colours) OTU numbers in sand dune successional stages. Data are averages of four deflation basins and three soil cores per basin. Ends of whiskers represent minimum and maximum values excluding outliers, and outliers are shown as separate points. Different letters indicate significant differences at p<0.05.

We sampled soil in each successional stage in four of the basins in July 2011. To take into account that plants and microbes vary at different spatial scales, the soil samples were collected in triplicate. The samples were taken from the depth of 0-10 cm with a 12.5 cm × 12.5 cm soil corer (72 samples in total: six successional stages × four basins × three soil cores per 1×1 m study plot). Soil was sieved with a 4-mm sieve and frozen at −20°C.

### Soil chemistry and temperature

Total C and N of soil organic matter, soil P, Ca, K, Na, Fe, Mn, Al, and Mg contents, and pH were measured as detailed in Methods S1. Soil temperature at the 10 cm depth was recorded every two hours with Hobo Temp External H08-002-02 loggers coupled with TMC6-HD soil temperature sensor during 2009-2015. We defined maximum temperature as the average temperature of the warmest month (July) and the minimum temperature as the average of the two coldest months (January and February).

### Phospholipid fatty acid (PLFA) analysis

Phospholipid fatty acids were extracted as described previously (Francini *et al*., 2014) from freeze-dried soil (15 g for sand and grass stages, 10 g for moss, 8 g for lichen, 5 g for heath and 3 g for forest stage corresponding to OM content). Briefly, lipids were extracted with chloroform:methanol:citrate buffer (Bligh and Dyer, 1:2:0.8 v/v/v), fractionated using silica columns (Bond Elut LRC, Varian), and phospholipids were methylated with mild alkaline methanolysis. The resulting fatty acid methyl esters were analyzed by gas chromatography and PLFAs identified based on a bacterial fatty acid standard mixture (Supelco, Bellefonte, PA, USA) and fatty acids from reference strains. For community analysis, data was expressed as peak area percentage of a single PLFA of the total area of the profile, and for biomass analysis as nmol of PLFAs per gram dry weight (gdw) of soil or per gram OM. The sum of PLFAs i15:0, a15:0, 15:0, i16:0, 16:1ω9, i17:0, a17:0/17:1ω8c (eluted in the same peak), 17:0, cyclo-17:0, 18:1ω7c/ω9t (eluted in the same peak) and cyclo-19:0 were used to represent bacterial biomass and PLFA 18:2ω6 fungal biomass. Sum of all these PLFAs was used to represent total microbial biomass. Biomass of selected bacterial groups was calculated by multiplying the total microbial biomass of a sample with the relative abundance of the bacterial group in the sample based on 16S rRNA gene sequencing.

### DNA extraction and PCR

DNA was extracted from 0.85 g (0.4 g for forest stage) soil with Powersoil DNA extraction kit (MoBio). Bead beating was carried out with a FastPrep instrument at 5.0 m s^-1^ for 40 s. DNA concentration was determined with Qubit fluorometer and Qubit dsDNA HS kit (Invitrogen).

Bacterial and fungal PCRs were carried out as two-step PCR where the second step introduced barcodes and adapters for Ion Torrent sequencing. The first step of bacterial 16S rRNA gene PCR (V1-V2 region) was carried out with primers 27F (5’-AGAGTTTGATCCTGGCTCAG-3’) and 338R (5’-TGCTGCCTCCCGTAGGAGT-3’).The first step of intergenic transcribed region (ITS) PCR for fungi was carried out with primers fITS7 (5’-GTGARTCATCGAATCTTTG-3’) and ITS4 (5’-TCCTCCGCTTATTGATATGC-3’) (Ihrmark *et al*., 2012), where the fITS7 primer contained an M13 adapter for the second step (Mäki *et al*., 2016).

The second step primers introduced barcodes and Ion Torrent adapters A and P1. For bacterial 2nd step PCR, the forward primer was adapter A-barcode-27F, and the reverse primer adapter P1-338R. For the fungal 2nd step PCR, the forward primer was adapter A-barcode-M13 adapter and the reverse primer adapter P1-ITS4. Details for PCR are given in Methods S1. After the second PCR, the duplicate products were pooled, purified with Agencourt AMPure XP purification system (Beckman Coulter), and quantified with Qubit fluorometer and Qubit dsDNA HS kit. Equimolar amounts (10 ng) of products were pooled for sequencing.

### Ion Torrent sequencing and sequence analysis

Pooled bacterial and fungal PCR products were sequenced after emulsion PCR with the Ion OneTouch system and Ion OT2 400 kit (Life Technologies) on Ion 314 chips with Ion PGM Sequencing 400 Kit (Life Technologies) according to the manufacturer’s instructions. The sequences were processed in mothur software package v. 1.38 (Schloss *et al*., 2009) following the relevant parts of MiSeq SOP protocol (https://mothur.org/wiki/MiSeq_SOP; accessed in January 2017; Kozich *et al*., 2013). Sequences were quality-filtered using average quality of 25 and a window size of 50 bases, a minimum sequence length of 200 bp, a maximum length of 400 bp for bacteria and 410 bp for fungi, and further settings maximum homopolymer length maxhomop=6, maximum number of ambigious bases maxambig=0, maximum number of differences to primer sequence pdiffs=1, and maximum number of differences to barcode sequence bdiffs=0. Bacterial sequences were aligned against Silva database v. 1.23 (Quast *et al*., 2013). Chimeras were detected with Uchime in de novo mode (Edgar *et al*., 2011) in mothur. Fungal ITS2 region was extracted and non-fungal sequences were removed with ITSx (Bengtsson-Palme *et al*., 2013). After quality filtering, alignment for bacteria, and removal of chimeras and non-target sequences, we had 121711 bacterial sequence reads and 279885 fungal reads. Reads were preclustered with setting diffs=2 (bacteria) or diffs=1 (fungi). The sequences were clustered into 97% operational taxonomic units (OTUs) using the average neighbor method for bacteria, and the vsearch algorithm for fungi, yielding 5242 bacterial and 1742 fungal OTUs after removal of singletons. Bacterial OTUs were classified against the Silva v. 1.23 database and fungi against the Unite database (Kõljalg *et al*., 2013). Data was subsampled to the read number of the sample with the lowest number of reads, which was 1055 reads for bacteria and 1605 reads for fungi. Shannon diversity index and coverage were calculated in mothur. Coverage was 76 ± 6% for bacteria and 97 ± 1% for fungi. The sequence data were submitted to NCBI under BioProject accession PRJNA471306.

### Statistical analyses

Chemistry and microbial biomass were compared among the successional stages using linear mixed models with R package nlme and function lme with successional stage as a fixed factor and deflation basin as a random blocking factor. Chemistry and biomass values except pH and Al were log transformed to normalize their distribution.

Bacterial and fungal sequence data were converted to relative abundances of OTUs and square root transformed to achieve Hellinger transformation that reduces the weight of OTUs with low counts and zeros. Only OTUs with 5 ≥ reads were included. Variation of community structure was visualized as non-metric multidimensional scaling (NMDS) plots with function metaMDS using the Bray-Curtis distance measure. Dispersion was visualized in the NMDS plots as ellipses based on standard deviation of sample scores using the ordiellipse function. Multivariate dispersion between the successional stages was tested with betadisper (Bray-Curtis distances). Permutational analysis of variance (PERMANOVA, Anderson, 2001; McArdle and Anderson, 2001) with the adonis2 function (Bray-Curtis distances) was used to compare community structure between the successional stages and in response to environmental variables. In the univariate model for successional stage, the stage was nested within deflation basin. The multivariate PERMANOVA model for environmental variables included OM-pH gradient and in addition Al and minimum soil temperature, which did not correlate with OM and pH (Pearson |r|<0.7) but differed with successional stage. The OM-pH gradient was defined with detrended correspondence analysis, and the first axis scores were used as a variable in PERMANOVA. Impact of the individual environmental variables was determined with marginal tests. Transitions between successional stages were quantified by calculating average distances of the successional stages from bare sand for bacterial and fungal community structure (Bray-Curtis distances), vegetation (Bray-Curtis distances of square root transformed data) and soil chemistry (Euclidean distances). Using the same distance measures, we carried out Procrustes analysis with the function procrustes to correlate microbial community structure with vegetation, and microbial community structure with soil chemistry (Lisboa *et al*., 2014). The analysis was based on the first four axes of NMDS analysis of bacterial community, fungal community, vegetation and soil chemistry (stress for all ordinations ≤0.07). The strength and the significance of the Procrustes correlation was tested with the function protest (Peres-Neto and Jackson, 2001). Residuals from the Procrustes analysis were used to compare the strength of the plant-microbe and soil chemistry-microbe correlation along the succession by comparing the successional stages with linear mixed models as above for soil chemistry. To separate the effects of vegetation and soil chemistry on the microbial community, we carried out partial Procrustes analysis where the variation due to soil chemistry was first partitioned out by multiple regression of vegetation and microbial data against soil chemistry, and the residuals were then used in Procrustes analysis (Peres-Neto and Jackson, 2001; Lisboa *et al*., 2014). All multivariate analyses were carried out with the vegan package (v.2.4.2, Oksanen *et al*., 2017) in R. Significance tests were based on 999 permutations.

We identified bacterial and fungal OTUs characteristic to specific successional stages or combinations of stages by indicator species analysis with the package indicspecies using the function multipatt and the association index IndVal.g which corrects for unequal site numbers (De Cáceres and Legendre, 2009). Only OTUs that occurred in the subsampled data at least 12 times, i.e., once in each sample of a successional stage, were included in the analyses. The combinations of stages included in the analyses were each successional stage on its own, all six stages together, all stages except sand, all stages except forest, all stages except sand and grass, all stages except heath and forest, sand+grass, moss+lichen, moss+grass, lichen+heath, heath+forest, sand+grass+moss, moss+lichen+heath, and lichen+heath+forest. Instead of using all possible combinations, we selected the combinations to limit the number of tests and to focus on the combinations most relevant for the aims of the study. OTUs were considered indicator OTUs if their indicator value was >0.3 and p <0.05. Significance was evaluated with a permutation test (999 permutations). P values were adjusted for multiple comparisons with function p.adjust (method “fdr”). Generalist OTUs present in all successional stages were defined at indicator values >0.5, because no p values are generated for taxa present in all sites. The identified indicator OTUs were grouped based on their taxonomic classification at order (bacteria) or class (fungi) level. Unclassified OTUs were included in these comparisons by re-clustering all reads assigned to unclassified bacteria or fungi in mothur at 75% sequence similarity to represent phylum level similarity for bacteria and class level similarity for fungi (Tedersoo *et al*., 2014).

To evaluate responses of bacterial and fungal indicator taxa to environmental variables, we used relative abundances of indicator OTUs grouped at order level (bacteria) and class level (fungi), including those unclassified 75% similarity groups that represented >0.4% of all unclassified OTUs. The tested environmental variables were those that best explained microbial community variation (OM, pH, Al for bacteria; OM, pH, minimum temperature for fungi). The analysis was carried out with the function traitglm in the package mvabund (v.3.13.1, Wang *et al*., 2012). The function fits fourth corner models to relate sites and their traits to species and species traits. When used without species trait data, it becomes a predictive model of responses of different OTUs to variation of environmental factors (Brown *et al*., 2014). The method used was glm1path, fitting a generalized linear model with LASSO penalty, which means that fourth corner coefficients of OTUs not decreasing Bayesian information criterion (BIC) are forced to 0, i.e. no response to environmental variation, to simplify the model (Brown *et al*., 2014).

All statistical analyses were carried out in R v.3.2.5. or v.3.4.0 (RCoreTeam, 2018).

## Results

### Soil chemistry and microbial biomass along Arctic sand dune succession

The increasing aboveground vegetation cover from bare sand to mountain birch forest was mirrored belowground in increasing soil OM content and decreasing pH (Fig. 1; Pearson correlation r=-0.78, p< 0.001). Organic matter content was low in the sand, grass, and moss stage soils (0.4±0.2%, mean±SD) but increased to 0.9±0.3% in the lichen stage and up to 4.5% ± 1.7% in the forest stage (Fig. 1b, Table S1). pH was highest in the grass stage (5.8 ± 0.2) and lowest in the forest stage (4.7 ± 0.1) (Fig. 1c). Soil contents of N, P, K, Ca, Mg, Fe and S increased with increasing OM (Table S1).

Microbial biomass measured as PLFAs per soil dry weight followed closely the OM-pH gradient (Fig. 1e, Fig. S1). The moss stage had similar OM content as bare sand, but its microbial biomass was higher than in the other sandy stages. When related to soil OM, microbial biomass in the moss stage reached a level similar to the lichen and vascular plant stages (Fig. 1d). Similarly, a shift to a higher proportion of fungal to bacterial biomass occurred in the moss stage and was maintained during the later successional stages (Fig. 1f). Overall, the proportion of fungi was highest in the heath stage.

### Microbial community structure

Bacterial and fungal community structure based on sequencing (Fig. 2) and PLFA (Fig. S2) followed the order of the vegetation stages on the OM-pH gradient and differed with successional stage (PERMANOVA bacteria r^2^=0.58 P<0.001, fungi r^2^=0.59 p<0.001). The moss stage grouped with the other sandy stages (sand, grass) with partly overlapping communities, whereas the lichen stage grouped with the vascular plant stages (heath, forest) (Fig. 2a, 2c). It should be noted that multivariate dispersion also differed between the successional stages (betadisper p=0.007 bacteria, p=0.001 fungi). The highest bacterial OTU richness (Fig. 1g) and Shannon diversity (Table S1) were detected in the heath stage, whereas fungal richness and diversity were highest though very variable in bare sand (Fig. 1g, Table S1). The soil parameter that best explained the community variation of bacteria and fungi was the OM-pH gradient, especially in the vegetated stages (Table S2).

**Figure 2.**
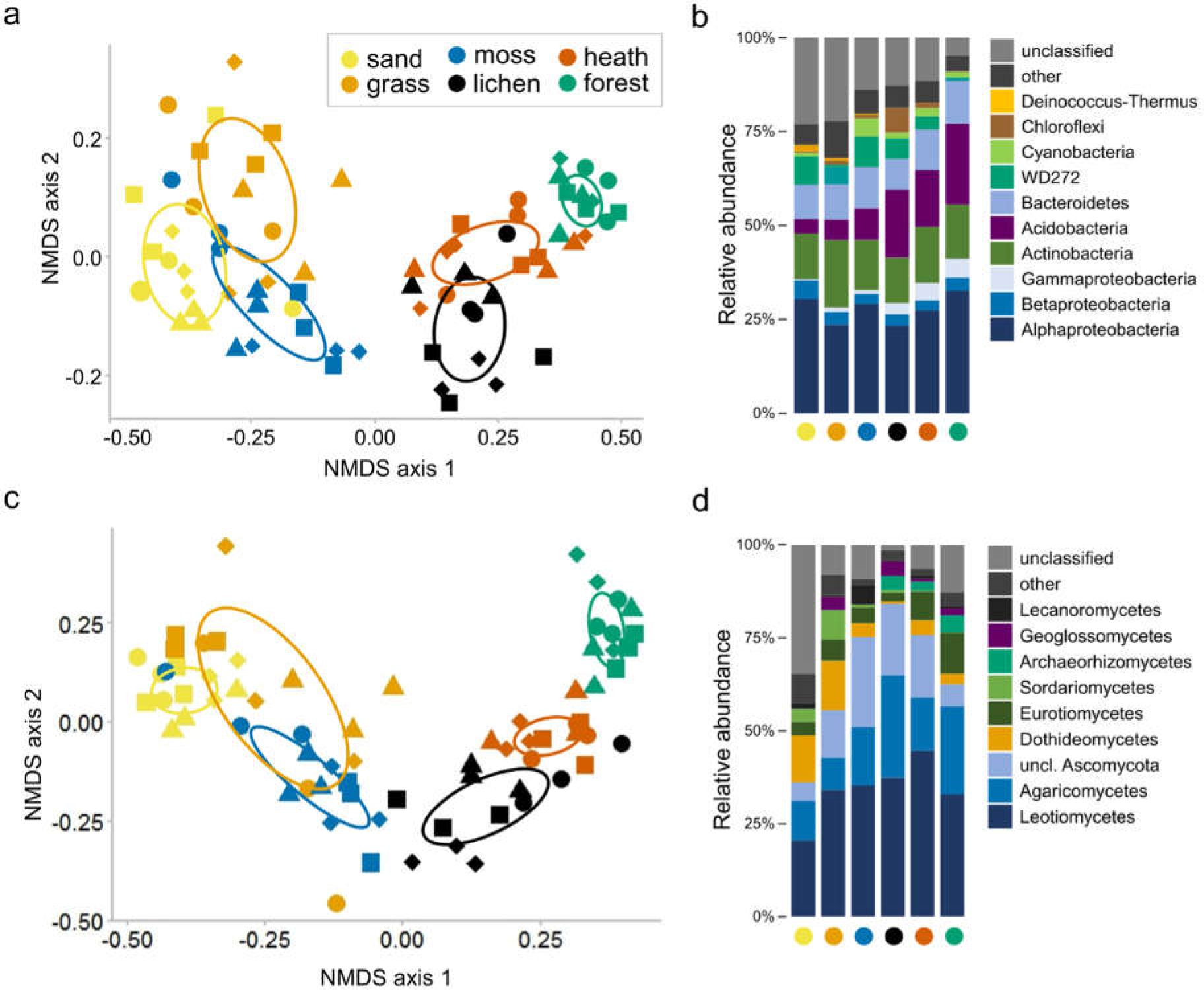
Community structure of bacteria (a) as a non-metric multidimensional scaling (NMDS) plot (stress=0.10) and (b) at phylum level (class level for Proteobacteria) taxonomical classification based on 16S rRNA gene amplicon sequencing, and community structure of fungi (c) as an NMDS plot (stress=0.13) and (d) at class-level taxonomical classification based on ITS2 region amplicon sequencing. Data points represent three soil cores from four deflation basins per sand dune successional stage. Shapes of the symbols represent separate deflation basins. In b and d, data are averages of four deflation basins and three soil cores per basin. For values of individual samples and the groups combined in ‘other’, see Fig. S4 and S7.

To compare successional shifts in bacteria, fungi, vegetation, and soil chemistry, we calculated distance measures from bare sand to the other successional stages (Fig. 3). The largest shift in bacterial and fungal community structure was detected from moss to lichen stage (Fig. 3). Soil chemistry changed most markedly from heath to forest. Vegetation showed increasingly larger shifts in later succession, with the largest shift from heath to forest (Fig. 3). To determine if the microbial communities correlated with vegetation and soil chemistry, we carried out Procrustes analysis (Fig. S3). The bacterial community correlated better with soil chemistry than vegetation (bacteria-vegetation r=0.71, bacteria-soil chemistry r=0.80, p=0.001 for both). The fungal community correlated tighter with vegetation than with soil chemistry (fungi-vegetation r=0.77, fungi-soil chemistry r=0.71, p=0.001 for both). When the variation due to soil chemistry was partitioned out, correlation with vegetation remained with lower correlation coefficients (bacteria-vegetation r=0.51, fungi-vegetation r=0.52, both p=0.001). We then used the Procrustes residuals to compare the strength of the relationship between the plant and microbial community along the succession. The strength of the vegetation-bacteria correlation did not vary with successional stage (Fig. 4). In contrast, vegetation determined fungal community composition more strongly in the heath stage than in the other stages (Fig. 4)

**Figure 3.**
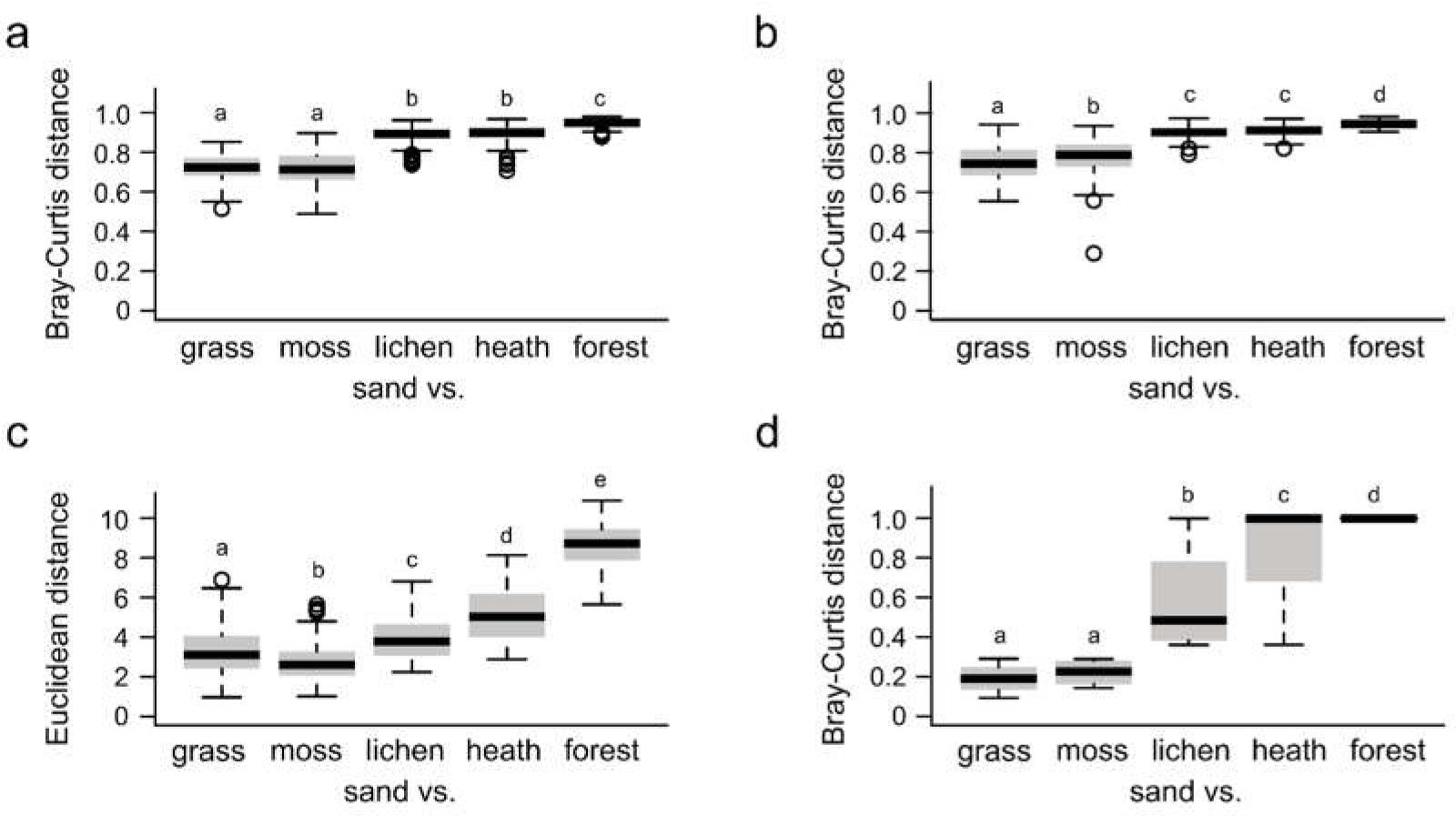
Average distances of successional stages from bare sand for bacterial, fungal, and plant communities and for soil chemistry. Different letters indicate significant differences at p<0.05. Ends of whiskers represent minimum and maximum values excluding outliers, and outliers are shown as separate points.

**Figure 4.**
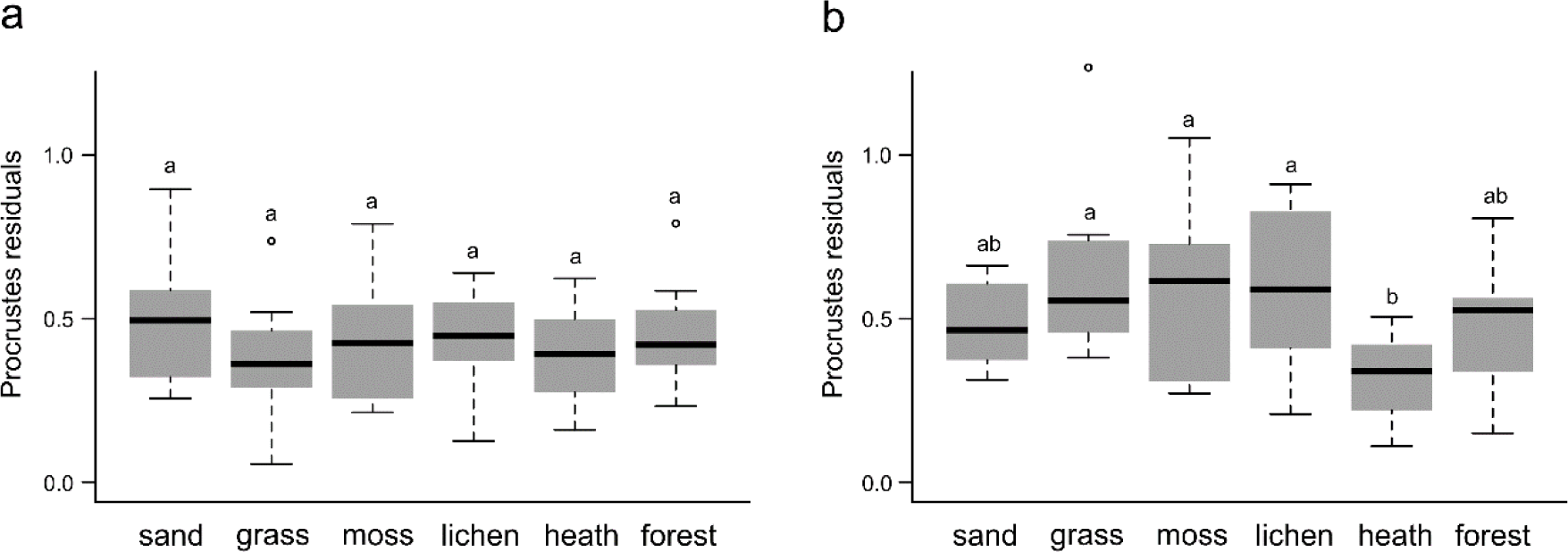
Strength of correlation of vegetation with (a) bacterial community and (b) fungal community determined as residuals of partial Procrustes analysis when variation due to soil chemistry was partitioned out. Low values of the residuals indicate stronger correlation. Different letters indicate significant differences between the sand dune successional stages at p<0.05.

### Bacterial taxonomic distribution with succession

Overall, the most abundant bacterial phyla or classes were Alphaproteobacteria, Actinobacteria, and Acidobacteria (Fig. 2b, Fig. S4). The most abundant bacterial OTU belonged to the candidate phylum WD272 (WPS-2). This phylum was the fourth or fifth most abundant phylum in the sandy and lichen stages (on average 4.6-8.1%), decreasing in the heath (3.5%) and forest (0.9%) (Fig. 2b, Fig. S4). When related to microbial biomass, the abundance of WD272 increased in the moss stage (Fig. S5). WD272 showed abundant indicator OTUs for the sandy stages but also less abundant forest-specific indicators (Fig. 5). Most indicator groups specific for the sandy stages were strongly associated with low OM content (Cytophagales, Deinococcales, Acidobacteria subgroup 4, Pseudonocardiales (Actinobacteria), Oligoflexales (Deltaproteobacteria), unclassified group 9) (Fig. 5). Cyanobacteria, three Chloroflexi groups and unclassified group 2 were indicators for the moss and lichen stages and associated with both low OM and low pH. Another sign that the moss stage was a key transition stage for bacterial community was that it showed unclassified clusters typical to both sandy and vegetated stages (Fig. S6a). Most indicator groups for the forest stage were indicators also for the other vegetated stages. For example, Acidobacteria subgroup 2 associated with low pH and subgroup 6 associated with high OM appeared in the moss and lichen stages and had indicators also in the heath and forest stages (Fig. 6). The overall abundance of Acidobacteria increased with decreasing pH (Pearson correlation r = −0.81, p<0.001).

**Figure 5.**
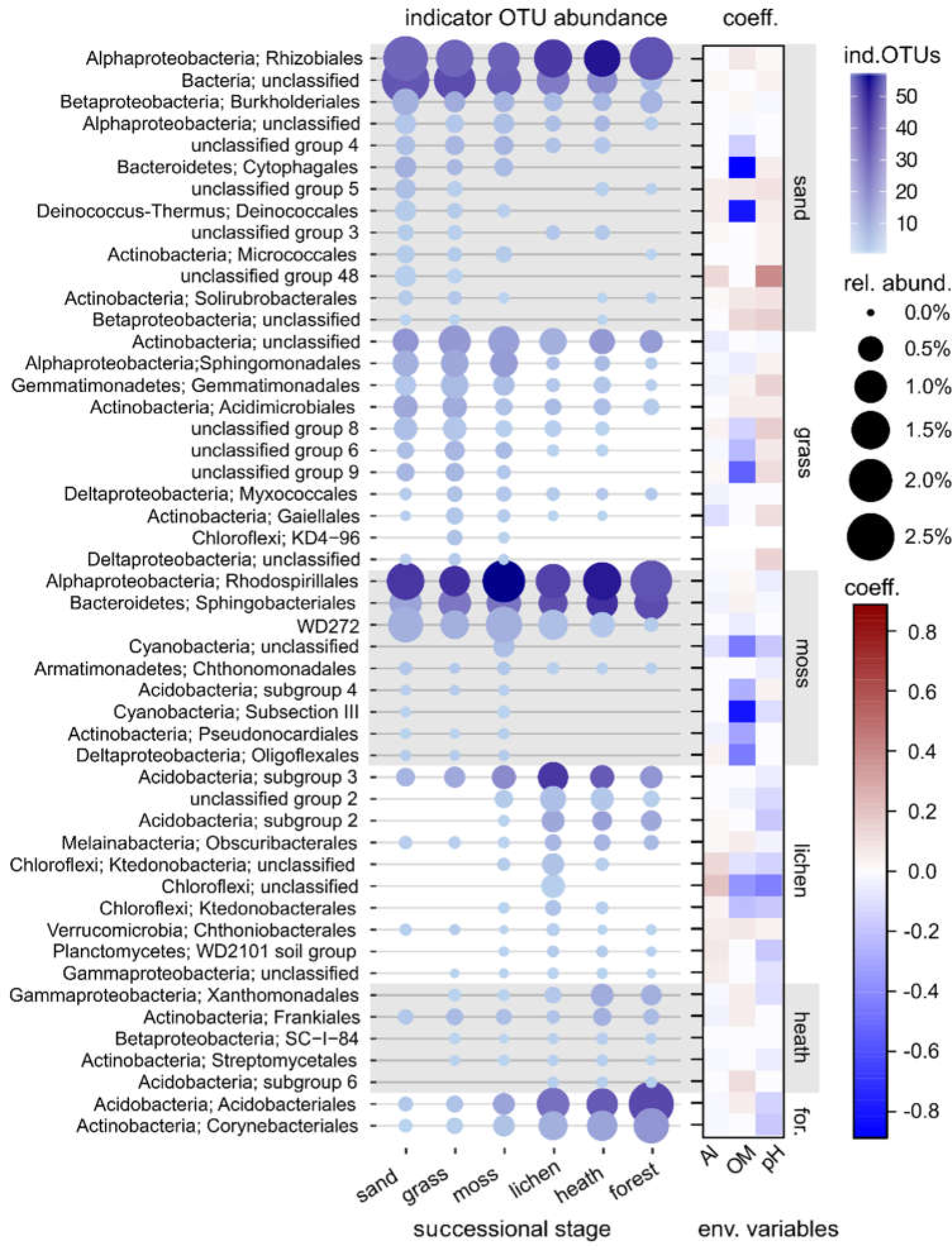
Bacterial indicator OTUs for successional stages grouped at order level. Grey shading shows where the successional stage changes, and each group is placed in the stage where it was the most abundant (vertical successional stage labels; for., forest). Only groups with total abundance of indicator OTUs >0.05% are shown. The heatmap on the right shows fourth corner coefficients (coeff.) describing the association of the indicator OTU groups with environmental variables (env. variables). Positive coefficient indicates that the microbial group occurred at high values of the environmental factor and negative coefficients indicate occurrence at low values of the environmental factor. Ind. OTUs, number of indicator OTUs; rel. abund., relative abundance; OM, organic matter.

**Figure 6.**
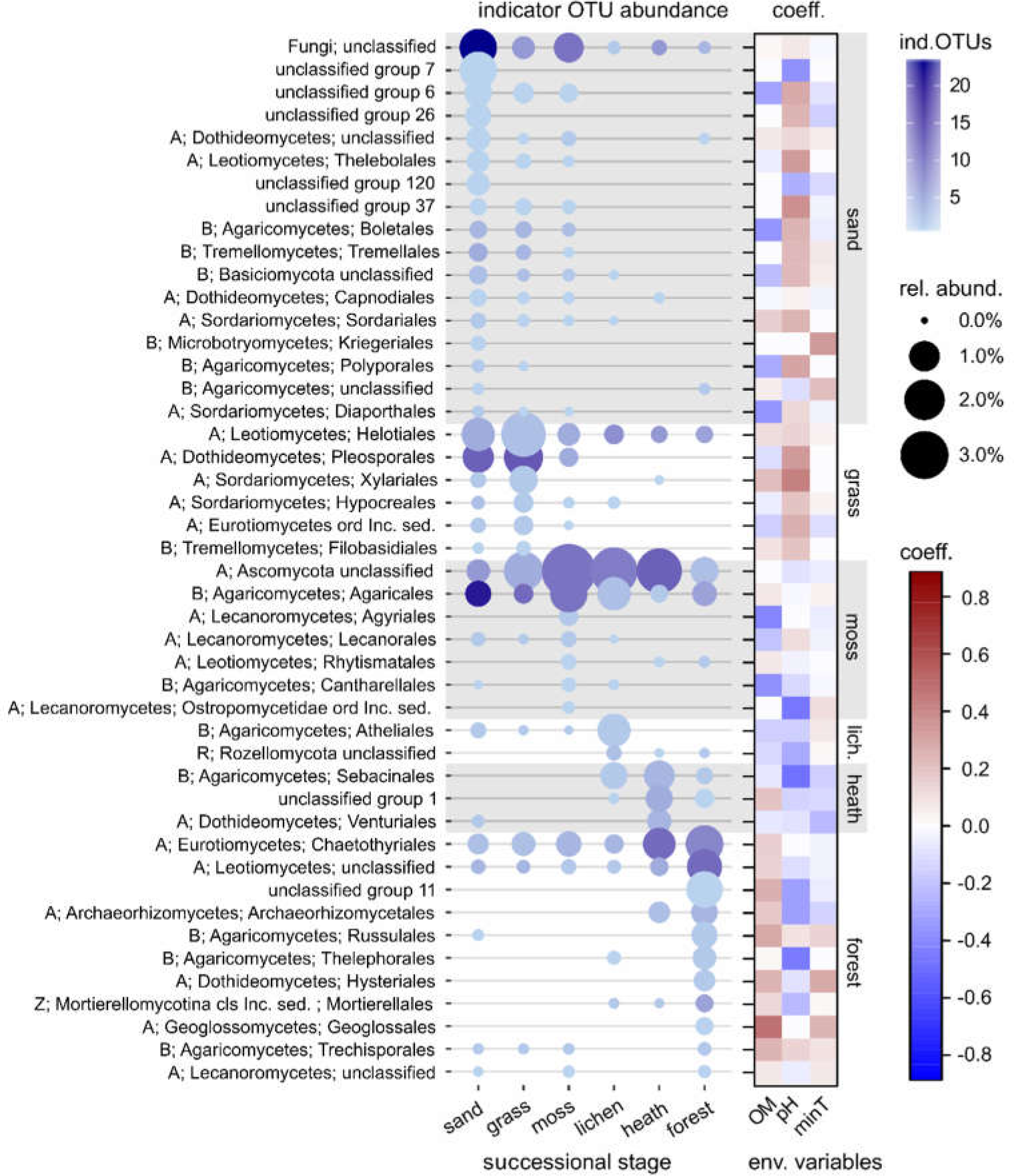
Fungal indicator OTUs for successional stages grouped at class level. Grey shading shows where successional stage changes, and each group is placed in the stage where it was the most abundant (vertical successional stage labels: lich., lichen). A, Ascomycota; B, Basidiomycota; R, Rozellomycota; Z, Zygomycota; cls, class; ord, order; Inc. sed., Incertae sedis. Only groups with total abundance of indicator OTUs >0.05% are shown. The heatmap on the right shows fourth corner coefficients (coeff.) describing the association of the indicator OTU groups with environmental variables (env. variables). Positive coefficient indicates that the microbial group occurred at high values of the environmental factor and negative coefficients indicate occurrence at low values of the environmental factor. Ind. OTUs, number of indicator OTUs; rel. abund., relative abundance; OM, organic matter, minT, minimum soil temperature.

### Fungal taxonomic distribution with succession

The fungi at class level were dominated by Ascomycetes, followed by Basidiomycetes, unclassified fungi, and smaller shares of Zygomycota, Rozellomycota, Chytridiomycota, and Glomeromycota (Fig. 2d, Fig. S7). The most abundant Ascomycota class was Leotiomycetes, followed by unclassified ascomycetes and Dothideomycetes. Bare sand had the highest relative abundance of unclassified fungi up to 35% (Fig. 2d, Fig. S7), and several unclassified clusters were specific to the sandy stages (Fig. S6b) and associated with high pH or low OM (Fig. 6). Other strong indicators of sandy stages included Helotiales and Thelebolales (Leotiomycetes), and Polysporales and Boletales (Agaricomycetes). Lecanoromycetes, containing lichenized fungi, was most abundant in the moss stage (Fig. 2d). Similarly to bacteria, the fungal groups that were the best indicators of the moss and lichen stages were associated with low OM and low pH or both, including Lecanoromycetes (Lecanorales, Agyriales, Ostropomycetidae), Agaricomycetes (Cantharellales, Atheliales, Agaricales), and Dothideomycetes (Pleosporales, Capnodiales) (Fig. 6). The most abundant fungal OTU overall and one of the fungal generalists (Table S3) was classified as *Pezoloma ericae* (Leotiomycetes; also known as *Rhizoscyphus ericae*). This OTU was particularly dominant in the heath (34±5% of reads). Its relative abundance was lowest in bare sand (2±1%) and the grass stage (12±14%), but in the moss and lichen stages it increased to 26±16%. In the forest, its relative abundance was lower than in the cryptogam-dominated stages (17±14%).

## Discussion

We hypothesized that the belowground and aboveground succession in Arctic sand dunes are not completely synchronous. Specifically, we hypothesized that the cryptogam-dominated stages of an Arctic sand dune succession involve key belowground components that facilitate transitions towards vascular vegetation. Supporting this hypothesis, the bacterial and fungal communities in the lichen stage soil resembled the communities in the later, vascular plant dominated stages. In the other cryptogam stage studied – the moss stage – the overall microbial communities overlapped with the earlier, sandy stages. Nevertheless, indicative of belowground transitions preceding aboveground succession, microbial biomass per OM and the proportion of fungi increased in the moss stage, and later stage microbial groups appeared despite the absence of their presumed host plants. The higher correlation of bacterial community with soil chemistry than with vegetation and the increase in the strength of plant-fungal interaction with vascular vegetation supported the hypothesis on the different influences of vegetation succession on bacteria and fungi.

The dune sand is a harsh habitat with low carbon and nitrogen availability. The initial microbial colonizers must be able to tolerate the oligotrophic conditions either by using the low levels of allocthonous organic matter (Hodkinson *et al*., 2002) or as autotrophs (Nemergut *et al*., 2007; Fierer *et al*., 2010). Comparison of biomass per OM with biomass per soil dry weight shows that the microbial biomass increased in the moss stage when OM potentially available for heterotrophs still remained low. A characteristic group for the early stages from bare sand to lichen was candidate phylum WD272, currently known as WPS-2 and recently named *Candidatus* Eremiobacterota, ‘desert bacterial phylum’, which are proposed to be autotrophs able to use atmospheric H_2_ and carbon monoxide (CO) as energy sources (Ji *et al*., 2017; Holland-Moritz *et al*., 2018). Another potential H_2_- and CO-oxidizing group in the moss and lichen stages were members of Chloroflexi classified as Ktedonobacteria (King and King, 2014; Islam *et al*., 2019). The increased occurrence of WD272 and Chloroflexi in the moss and lichen stages suggests that they may form a basis of biomass accumulation in the Arctic sand dune succession. These microbes may therefore facilitate development of cryptogam cover and heterotrophic microbial community (Connell and Slatyer, 1977; Brooker *et al*., 2008).

Many moss stage microbial groups that were rare or absent in the earlier sandy stages are linked to biological soil crusts. Biocrusts are consortia of microbes, mosses, and lichens that are recognized as the major stabilizing factor in early successional soils worldwide (Weber *et al*., 2016). The rhizoids of *Polytrichum* mosses themselves stabilize soil, facilitate N_2_ fixation, and provide OM to the soil (Bowden, 1991). The moss stage showed the highest proportion of Cyanobacteria, known as N_2_-fixing early colonizers and components of biocrusts. Another distinctive indicator for the cryptogam stages were Chloroflexi, which in dryland and grassland ecosystems occur in moss and lichen biocrusts and in sub-crust soil (Kuske *et al*., 2012; Navarro-Noya *et al*., 2014; Maier *et al*., 2018). In a temperate inland dune succession, a thick *P. piliferum* cover increased microbial biomass compared to bare sand (Sparrius and Kooijman, 2013). In our ecosystem, *P. piliferum* biomass was small in the moss stage but nonetheless affected the microbial biomass and communities. Therefore *P. piliferum* appeared to have a disproportionate effect on the ecosystem level, suggesting it is a keystone plant species in Arctic sand dunes. The mechanism could be soil stabilization by rhizoids and moss-associated, soil crust-forming microbes. Traditionally, grasses are credited with the initial physical stabilization of sand in dune succession (e.g. Olson, 1958; Ellenberg, 2009). Inspecting the soil microbial community structure, it seems that in these Arctic dunes the grass *Deschampsia flexuosa* may not drive ecosystem succession. Based on our results, we propose that *P*. *piliferum* could participate in the physical stabilization of Arctic inland dunes and direct succession towards vascular vegetation. Verifying this proposition requires further studies on *P. piliferum* with emphasis on the role of rhizoids.

Indicative of nonaligned belowground vs. aboveground succession, lichen-forming fungi (Lecanoromycetes, Ascomycetes) were detected in the moss stage soil before the succession stage where they formed lichens. Furthermore, the relative abundance of the dominant fungal OTU *Pezoloma ericae* increased in the moss stage to levels similar to those in the later stages. We interpret this increase as a possible sign of fungal community shift at the moss stage towards a status where vascular plant establishment is possible. *Pezoloma ericae* forms mycorrhiza with ericoid plants such as the dominant heath species *Empetrum nigrum* (Read, 1991; Smith and Read, 2008). However, no known host plants of *P. ericae* were detected in the moss stage. In addition to ericoid mycorrhiza, *P. ericae* forms mycorrhiza-like associations in liverwort rhizoids (Pressel *et al*., 2010; Kowal *et al*., 2018), and it has been found as an Antarctic moss endophyte (Zhang *et al*., 2013). However, no liverworts were visible among the moss at our site, and no association of *P. ericae* with *Polytrichum* rhizoids is known.

Supporting the hypothesis of different controls, bacteria and fungi showed distinct richness and biomass patterns along the succession. The proportion of fungi in the PLFA data, assumed to represent the living biomass (Zhang *et al*., 2019), increased already in the moss stage. Bacterial richness peaked in the heath and fungal richness in the bare sand, and neither correlated with ecosystem productivity or community maturity defined as soil OM or plant richness. In general, richness and diversity link to ecosystem functions through the complex interaction between diversity and ecosystem stability and productivity (Loreau and de Mazancourt, 2013). Plant richness peaks in the middle or late succession (Bonet and Pausas, 2004; Peyrat and Fichtner, 2011; Prach *et al*., 2014). For microbes, few general patterns of richness or diversity with primary succession have emerged. In very young soils, richness is often low and then increases and stabilizes or even decreases in later stages (Jackson, 2003; Nemergut *et al*., 2007; Fierer *et al*., 2010; Williams *et al*., 2013; Jiang *et al*., 2018). Primary succession gradients where both bacteria and fungi have been studied show conflicting patterns with each other but generally agree on different trajectories for bacterial and fungal richness (Brown and Jumpponen, 2014; Cutler, *et al*., 2014; Dini-Andreote, *et al*. 2014; Dini-Andreote, *et al*. 2016; Jiang, *et al*. 2018). In our case, fungal richness remained rather stable under vegetation covered stages, while bacterial richness had a unimodal response. What can be concluded from above and the present study is that fungi and bacteria do not follow the same trajectories during succession and are not equally shaped by the same factors.

The low bacterial richness in the bare sand stage can be related to the oligotrophic conditions (i.e. low environmental resource availability); richness already increased with the scattered presence of the grass *D. flexuosa*. Pioneering plants can influence bulk soil when plants are of sufficient size (Tscherko *et al*., 2004; Edwards *et al*., 2006; Miniaci *et al*., 2007). The high fungal richness in the bare sand together with the low fungal biomass, however, is unpredicted by theories that species richness should increase with the increase in resources and environmental heterogeneity (Stein *et al*., 2014; Cline *et al*., 2018). Some of the fungal richness in bare soil may represent dormant fungi, spores or hostless biotrophic fungi (Jumpponen, 2003; Rime *et al*., 2015). On the other hand, our observation that the unclassified fungi can be grouped into class level clusters that occurred consistently in replicate samples and across the sandy successional stages also hints that the fungal community in bare sand may be in one way or other selected or specialized.

The high bacterial richness in the heath coincided with the highest fungal biomass and highest occurrence of *P. ericae*, presumably forming mycorrhizae with the dominant ericoid plant *E. nigrum*. We suppose that the high bacterial richness is promoted by resources and niches provided by fungal mycelium and *Empetrum* litter, despite its high phenolic content and proposed adverse properties on microbial activity (Wardle *et al*., 1998; Gallet *et al*., 1999). In the forest, where the highest microbial richness would be expected if richness correlated positively with OM, plant species richness, and productivity, bacterial diversity was lower than in the heath, possibly due to the environmental filtering effect of the low pH in the forest soil (Tripathi *et al*., 2018). The varied responses of bacterial and fungal richness and diversity to primary succession highlight the need to cover successional gradients with different parent material, vegetation and soil chemistry to understand microbial community patterns along succession.

Effects of soil chemistry on microbial communities can override biotic factors such as plant richness (Chen *et al*., 2016; Jiang *et al*., 2018), and the role of vegetation is emphasized in successional gradients where soil chemistry does not change markedly (Knelman *et al*., 2012; Cutler *et al*., 2014). Soil chemistry, vegetation and microbes are strongly intertwined in succession, and the present gradient from bare sand to mountain birch forest represents the common pattern where organic matter increases and the soil acidifies with increasing plant cover (Odum, 1969; Merilä *et al*., 2010). However, the largest microbial community shifts did not align with the largest shifts in soil chemistry or vegetation. No measured soil chemistry variable changed in parallel with the large bacterial community shift between moss and lichen stages. Both cryptogam stages showed indicator microbial groups associated with low OM and low pH. These results suggest that the microbial shift from moss to lichen stage is due to unmeasured effects of the lichen crust formation. Such factors could be better long-term moisture retention by the lichen crust than the sparse mosses in the moss stage, microenvironments provided by lichen crumbs in the belowground soil (de los Ríos *et al*., 2011) or the different chemical composition and secondary metabolites of mosses versus lichens.

### Conclusions

This study on inland sand dunes illuminates bacterial and fungal succession in relation to vegetation and soil chemistry in a less studied Arctic soil ecosystem. Successional dynamics may not be linear nor deterministic, which makes the space-for-time substitution in mosaics of repeated plant communities that are thought to represent chronosequences challenging (Walker *et al*., 2010). In our case, the soil organic matter helps link vegetation development in the lichen, heath and forest with time. However, the moss and grass stages are equally devoid of soil organic matter. The microbial community composition in the grass stage suggests that this stage is not well connected to the progressive successional pathway from bare sand to mountain birch forest. Our results highlight the importance of considering not only vascular plants but also cryptogams, i.e. mosses and lichens and particularly the potential dune keystone species *Polytrichum piliferum*, when defining the plant-microbe interactions in succession. Taken together, the results indicate that the belowground transition towards the later stages starts before the establishment of vascular vegetation. This means that the belowground succession may have a conditioning effect on the vascular plant succession in the early primary succession. This work complements evidence that bacterial and fungal richness are not necessarily related to soil ecosystem productivity or change in plant species richness (Waldrop *et al*., 2006; Porazinska *et al*., 2018; Delgado-Baquerizo and Eldridge, 2019). Soil chemistry and the effects of plants on microbial resources and niches seem to be important factors driving succession. However, the key transitions in soil chemistry, vegetation and microbial community did not align, suggesting additional drivers such as competition between microbial taxa. Bacterial and fungal indicators identified in this work, including novel unclassified diversity, can signpost ecosystem transitions towards vascular plant cover. Understanding the drivers of the shift from bare to vegetated stage is crucial for restoration of vegetation on cold-climate soils suffering from erosion and vegetation loss under changing environment.

## Acknowledgements

We thank Helena Jauhiainen, Riitta Nissinen and Manoj Kumar for help with sampling and DNA extraction, Anita Mäki for assistance with sequencing, and Outi Manninen for the vegetation inventory. This work was funded by Maj and Tor Nessling Foundation and the Academy of Finland (project 287545 to MMK).

## Author contributions

MMK conceived the research and led the field work, HJ and MM conducted PLFA analyses, HJ conducted DNA work, analyzed all the data and performed the statistical analyses, MT provided expertise and resources for Ion Torrent sequencing, HJ and MMK wrote the manuscript, and all authors commented on the manuscript.

