## Supplementary materials for "Cryptogams signify key transition of bacteria and fungi in Arctic sand dune succession"

**Methods S1:** Soil chemistry and temperature; Bacterial and fungal PCR

**Table S1:** Environmental parameters and microbial characteristics of successional stages

**Table S2:** Environmental variables explaining microbial community variation

**Table S3:** Bacterial and fungal generalist OTUs based on indicator species analysis.

**Figure S1:** Bacterial and fungal biomass

**Figure S2:** NMDS plot of PLFA data

**Figure S3:** Strength of correlation of microbial community vs. vegetation and chemistry

**Figure S4:** Phylum level community composition of bacteria with successional stage

**Figure S5:** Biomass estimates of selected bacteria with successional stage

**Figure S6:** Unclassified bacterial phylum level groups and fungal class level groups

**Figure S7:** Class-level community composition of fungi with successional stage

**Methods S1**

**Soil chemistry and temperature**

Total C and N of soil organic matter were analysed on a LECO CHN 1000 elemental analyser (LECO Corporation, USA). For analyzing soil P, Ca, K, Na, Fe, Mn, Al, and Mg contents, soils were extracted with acid (pH 4.65) 1 M ammonium acetate (15 ml soil sample in 150 ml extraction solution) and the elements were determined with an ICP emission spectrometer iCAP 6500 DUO 2011 (Thermo Scientific, United Kingdom). The dry matter content of the soil was determined by drying the samples (105 ºC) and organic matter (OM) content by loss on ignition (550 ºC) using LECO TGA-701 analyzer (LECO Corporation, USA). pH was determined in 1:2 (w:v) soil:water suspensions (10 g soil and 20 ml $H_2O$). Soil temperature at the 10 cm depth was recorded every two hours with Hobo Temp External H08-002-02 loggers coupled with TMC6-HD soil temperature sensor during 2009-2015. We defined maximum temperature as the average temperature of the warmest month (July) and the minimum temperature as the average of the two coldest months (January and February).

**Bacterial and fungal PCR**

Bacterial and fungal PCRs were carried out as two-step PCR where the second step introduced barcodes and adapters for Ion Torrent sequencing. The first step of bacterial 16S rRNA gene PCR (V1-V2 region) was carried out with primers 27F (5'-AGAGTTTGATCCTGGCTCAG-3') and 338R (5'-TGCTGCCTCCCGTAGGAGT-3'). The 25-µl reactions were carried out in duplicate and contained 1 × buffer, 200 µM dNTPs, 0.4 µM each primer, 0.4µg $\mu l^{-1}$ BSA, 0.75 U DreamTaq DNA polymerase (Thermo Fisher Scientific), and 1-2 µl of template DNA (depending on concentration). The cycling conditions were initial denaturation at 95 °C for 3 min, followed by 30 cycles of 95 °C for 30 s, 53 °C for 30 s, and 72 °C for 45 s, and a final extension of 72 °C for 7 min. The first step of intergenic transcribed region (ITS) PCR for fungi was carried out with primers fITS7 (5'-GTGARTCATCGAATCTTTG-3') and ITS4 (5'-TCCTCCGCTTATTGATATGC-3') (Ihrmark *et al.*, 2012), where the fITS7 primer contained an M13 adapter for the second step (Mäki *et al.*, 2016). Reaction composition was the same as in bacterial PCR but including additional $MgCl_2$ to total concentration of 3 mM. The cycling conditions were initial denaturation at 94 °C for 3 min, followed by 30 cycles of 94 °C for 30 s, 55 °C for 30 s, and 72 °C for 30 s, and a final extension of 72 °C for 7 min.

The second step primers introduced barcodes and Ion Torrent adapters A and P1. For bacterial 2nd step PCR, the forward primer was adapter A-barcode-27F, and the reverse primer adapter P1-338R. For the fungal 2nd step PCR, the forward primer was adapter A-barcode-M13 adapter and the reverse primer adapter P1-ITS4. Second step PCR reactions (25 µl) for both bacteria and fungi contained 1 × buffer, 200 µM dNTPs, 0.3 µM each primer, 0.5 U DreamTaq DNA polymerase, and 1 µl (bacteria) or 2 µl (fungi) of 1:10-diluted first step PCR product as template. The cycling conditions were as in the first step PCR but with 7 cycles. After the second PCR, the duplicate products were pooled, purified with Agencourt AMPure XP purification system (Beckman Coulter), and quantified with Qubit fluorometer and Qubit dsDNA HS kit. Equimolar amounts (10 ng) of products were pooled for sequencing. The fungal product pool was size-separated in precast 1% Pippin Prep gel (Sage Science) to remove oversized products > 400 bp, which occurred in some of the samples. These products were cloned and sequenced to confirm they were not of fungal origin (data not shown).

**Table S1.** Vegetation, soil chemistry, environmental parameters and microbial characteristics of successional stages in sand dune deflation basins. Data are mean values ± SD of six deflation basins (vegetation, soil temperature) or four deflation basins (chemistry, microbial data) and three replicates per deflation basin.

| successional stage | sand | grass | moss | lichen | heath | forest |
|---|---|---|---|---|---|---|
| vegetation | - | Deschampsia flexuosa | Polytrichum piliferum | Stereocaulon sp., Racomitrium ericoides, P. piliferum, Cladonia sp. | Empetrum nigrum | E. nigrum, Vaccinium vitis-idaea, Pleurozium schreberi, D. flexuosa, Linnea borealis |
| sand cover (%) | 100 ± 0 | 95 ± 3 | 92 ± 8 | 52 ± 36 | 39 ± 48 | 0 ± 0 |
| organic matter (%) | 0.4 ± 0.2 | 0.3 ± 0.1 | 0.5 ± 0.2 | 0.9 ± 0.3 | 1.8 ± 1.0 | 4.5 ± 1.7 |
| pH | 5.6 ± 0.2 | 5.8 ± 0.2 | 5.3 ± 0.2 | 5.2 ± 0.1 | 5.2 ± 0.2 | 4.7 ± 0.1 |
| snow cover (cm)[a] | 95±6 | 86±6 | 80±4 | 85±3 | 46±5 | 84±5 |
| moisture (%) | 0.16±0.02 | 0.17±0.04 | 0.17±0.02 | 0.25±0.05 | 0.36±0.11 | 0.51±0.14 |
| N (%) | 0.021 ± 0.002 | 0.022 ± 0.001 | 0.025 ± 0.003 | 0.036 ± 0.005 | 0.047 ± 0.014 | 0.088 ± 0.024 |
| P (mg/kg) | 2.4 ± 0.6 | 1.9 ± 0.7 | 3.1 ± 1.3 | 4.3 ± 1.2 | 6.7 ± 2.8 | 14.1 ± 3.2 |
| Al (mg/kg) | 118 ± 42 | 64 ± 28 | 91 ± 32 | 128 ± 42 | 94 ± 19 | 70 ± 27 |
| Ca (mg/kg) | 42 ± 64 | 84 ± 57 | 40 ± 33 | 50 ± 59 | 132 ± 90 | 269 ± 104 |
| Mg (mg/kg) | 9 ± 15 | 19 ± 16 | 8 ± 6 | 9 ± 9 | 27 ± 20 | 103 ± 30 |
| Fe (mg/kg) | 6.1 ± 1.6 | 6.7 ± 1.0 | 6.9 ± 1.5 | 9.8 ± 3.0 | 6.1 ± 1.8 | 24.9 ± 15.1 |
| S (mg/kg) | 2.4 ± 0.5 | 1.8 ± 0.6 | 2.8 ± 0.7 | 4.0 ± 0.8 | 4.8 ± 1.5 | 8.8 ± 3.3 |
| microbial biomass (nmol/g soil dw)[b] | 1.69±0.8 | 2.18±1.5 | 9.16±5.9 | 17.9±5.2 | 43.7±25.7 | 98.7±33.9 |
| fungal:bacterial biomass[b] | 0.04±0.02 | 0.04±0.04 | 0.24±0.13 | 0.18±0.07 | 0.48±0.14 | 0.29±0.10 |
| minimum soil temperature (°C)[c] | -3.8 ± 0.6 | -3.9 ± 0.6 | -3.5 ± 0.3 | -2.8 ± 0.5 | -6.2 ± 1.5 | -2.2 ± 1.1 |
| maximum soil temperature (°C)[d] | 15.3 ± 1.1 | 15.3 ± 0.3 | 15.3 ± 0.2 | 15.0 ± 0.8 | 14.6 ± 0.2 | 10.7 ± 0.8 |
| bacterial diversity (Shannon) [e] | 5.0 ± 0.3 | 5.3 ± 0.3 | 5.4 ± 0.1 | 5.5 ± 0.2 | 5.8 ± 0.1 | 5.3 ± 0.2 |
| fungal diversity (Shannon)[f] | 3.6 ± 0.7 | 2.8 ± 0.8 | 2.7 ± 0.6 | 2.4 ± 0.2 | 2.8 ± 0.8 | 2.9 ± 0.5 |

[a] maximum snow depth (cm) measured in late March-early April, mean values ± SE in years 2016, 2017, 2018
[b] based on phospholipid fatty acid (PLFA) analysis
[c] average temperature in January-February
[d] average temperature in July
[e] based on bacterial 16S rRNA gene sequencing
[f] based on fungal ITS2 region sequencing

**Table S2** Environmental variables explaining microbial community variation in sand dune successional stages (% variation explained in PERMANOVA (p), p<0.05 in bold).

| group | stage | OM-pH[a] | Al | Tmin[d] | all |
|---|---|---|---|---|---|
| bacteria | all stages | **21.4 (0.001)** | **3.8 (0.002)** | **2.5 (0.007)** | **28.7 (0.001)** |
| | sandy[b] | **8.3 (0.001)** | **8.8 (0.005)** | 3.8 (0.055) | **21.9 (0.001)** |
| | vegetated[c] | **10.6 (0.001)** | **5.1 (0.003)** | **5.3 (0.006)** | **31.0 (0.001)** |
| fungi | all stages | **16.2(0.001)** | **2.7 (0.006)** | **3.9 (0.001)** | **23.5 (0.001)** |
| | sandy[b] | **5.2 (0.018)** | **5.9 (0.006)** | 5.1 (0.230) | **16.3 (0.001)** |
| | vegetated[c] | **10.2 (0.001)** | **4.5 (0.003)** | **8.5 (0.001)** | **32.4 (0.001)** |

[a] gradient of OM and pH from detrended correspondence analysis

[b] sand, grass, moss

[c] lichen, heath, forest

[d] minimum temperature, average temperature in January-February

**Table S3.** Bacterial and fungal generalist OTUs based on indicator species analysis.

| kingdom | number of OTUs | taxonomic classification |
|---|---|---|
| bacteria | 1 | Proteobacteria, Alphaproteobacteria, Rhizobiales, unclassified |
| | 4 | Proteobacteria, Alphaproteobacteria, Rhizobiales, Xanthobacteraceae |
| | 3 | Proteobacteria, Alphaproteobacteria, Rhodospirillales, Acetobacteraceae |
| | 1 | Proteobacteria, Betaproteobacteria, Burkholderiales, unclassified |
| | 1 | Proteobacteria, Betaproteobacteria, Burkholderiales, Comamonadaceae |
| | 1 | Proteobacteria, Gammaproteobacteria, Xanthomonadales, Incertae Sedis |
| | 1 | Actinobacteria, Thermoleophilia, Solirubrobacterales, unclassified |
| | 2 | Acidobacteria, Acidobacteriales, Acidobacteriaceae |
| | 1 | Acidobacteria, Subgroup 2 |
| fungi | 1 | Ascomycota, Leotiomycetes, Leotiales, Leotiaceae (Pezoloma ericae) |
| | 1 | Ascomycota, Leotiomycetes, Helotiales, Hyaloscyphaceae |
| | 1 | Ascomycota, Leotiomycetes, Helotiales, unclassified |
| | 1 | Ascomycota, Dothideomycetes, Venturiales, Venturiaceae |
| | 1 | Ascomycota, Lecanoromycetes, unclassified |
| | 1 | Ascomycota, Pezizomycetes, Pezizales, Sarcosomataceae (Pseudoplectania nigrella) |
| | 1 | Ascomycota, unclassified |

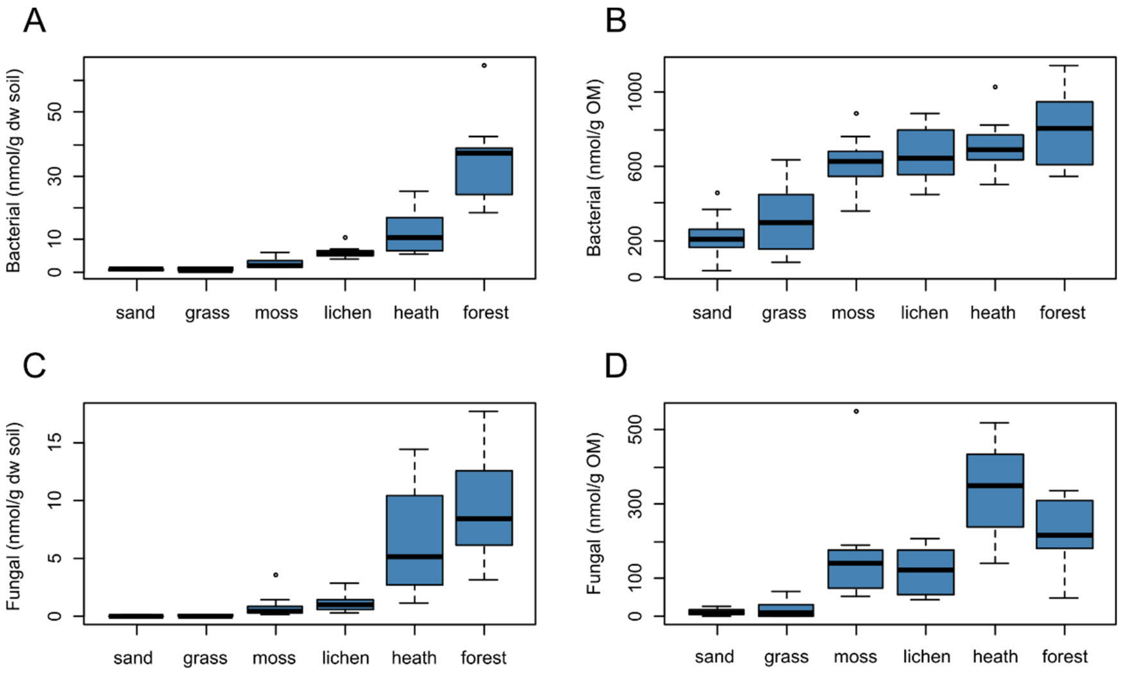


**Figure S1** Bacterial (A, B) and fungal (C, D) biomass based on phospholipid fatty acid analysis (PLFA) per soil dry weight (dw) (A, C) and soil organic matter (OM (B, D) with sand dune succession stage. Data are averages of four deflation basins and three replicates per basin ±SD. Note different scales on the y axis.

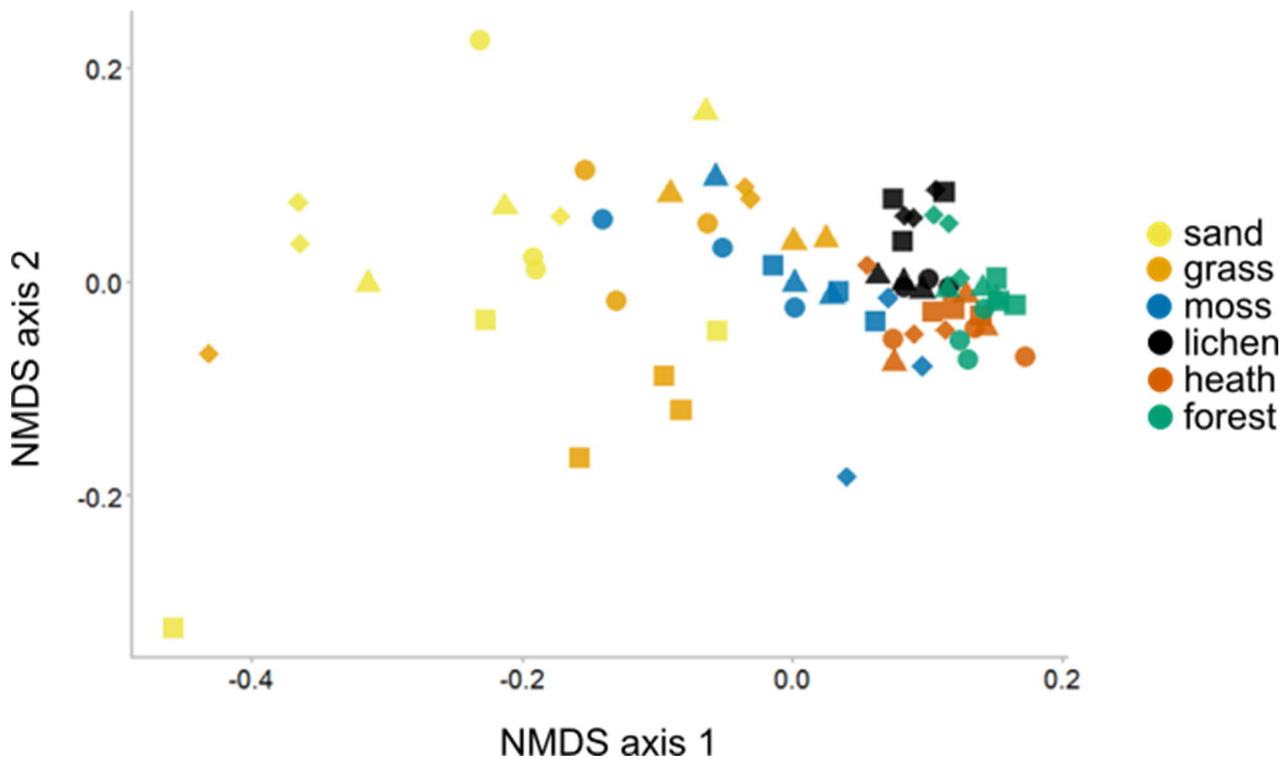


**Figure S2** Non-metric multidimensional scaling (NMDS) plot of phospholipid fatty acid (PLFA) data of sand dune successional stages (stress= 0.11). Shapes of the symbols refer to four different deflation basins.

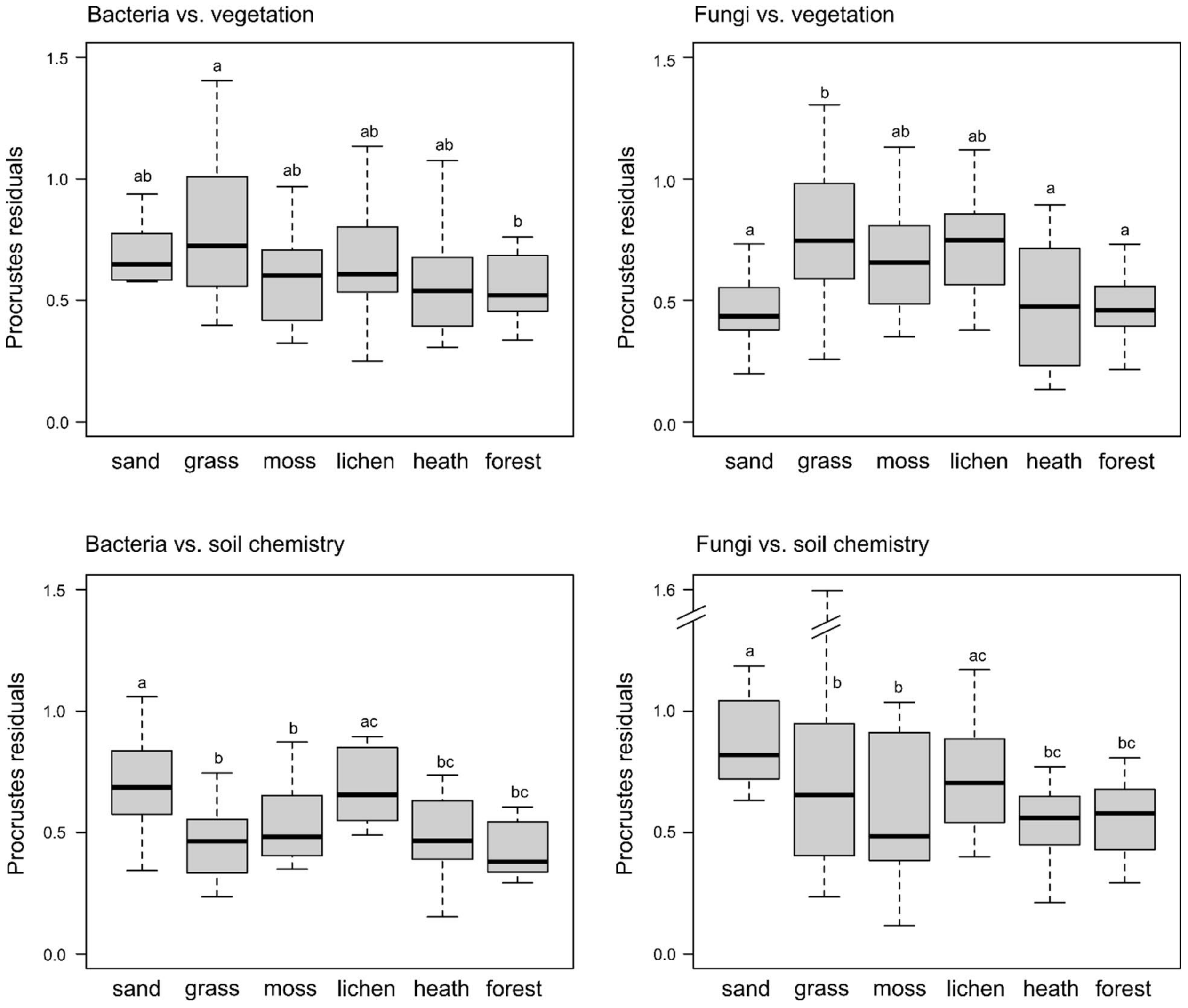


**Figure S3** Strength of correlation of bacterial community vs. vegetation, fungal community vs. vegetation, bacterial community vs. soil chemistry, and fungal community vs. soil chemistry with sand dune successional stages. Strength of correlation was determined as residuals Procrustes analysis, and low values of the residuals indicate stronger correlation. Different letters indicate significant differences between the sand dune successional stages at $p<0.05$.

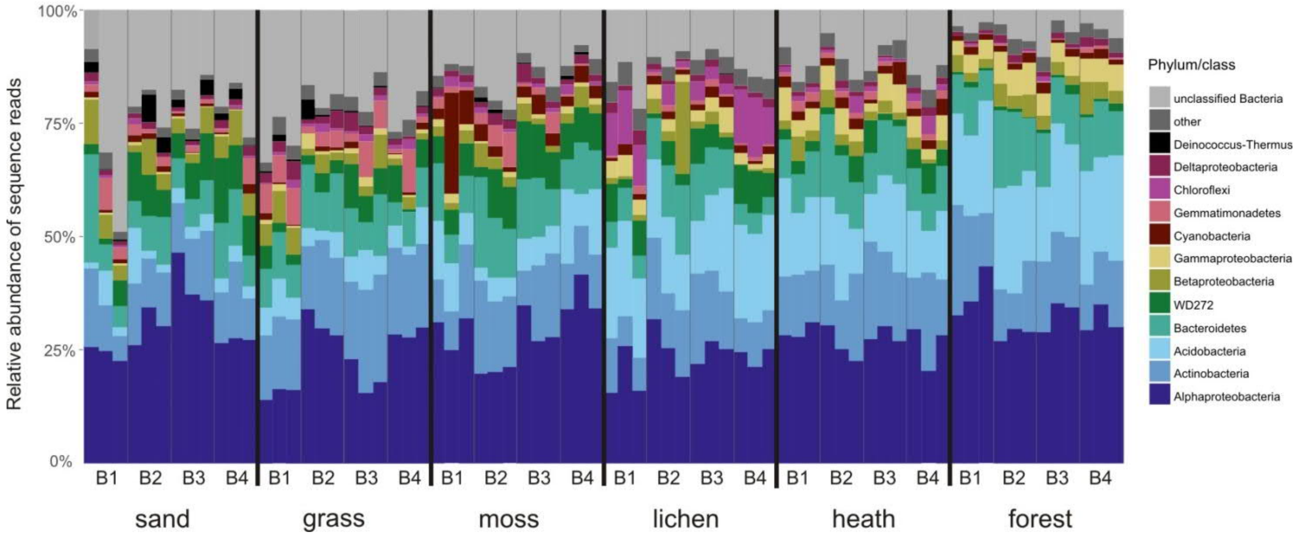


**Figure S4** Phylum level (class level for Proteobacteria) community composition of bacteria based on 16S rRNA gene sequencing of six sand dune successional stages. B1-B4 indicate four deflation basins, and three columns within a deflation basin are replicate soil samples. 'Other' includes Armatimonadetes, Verrucomicrobia, Planctomycetes, Saccharibacteria, unclassified Proteobacteria, Gracilibacteria, TM6, Firmicutes, Nitrospirae, Elusimicrobia, Parcubacteria, WCHB1-60, and Hydrogenedentes.

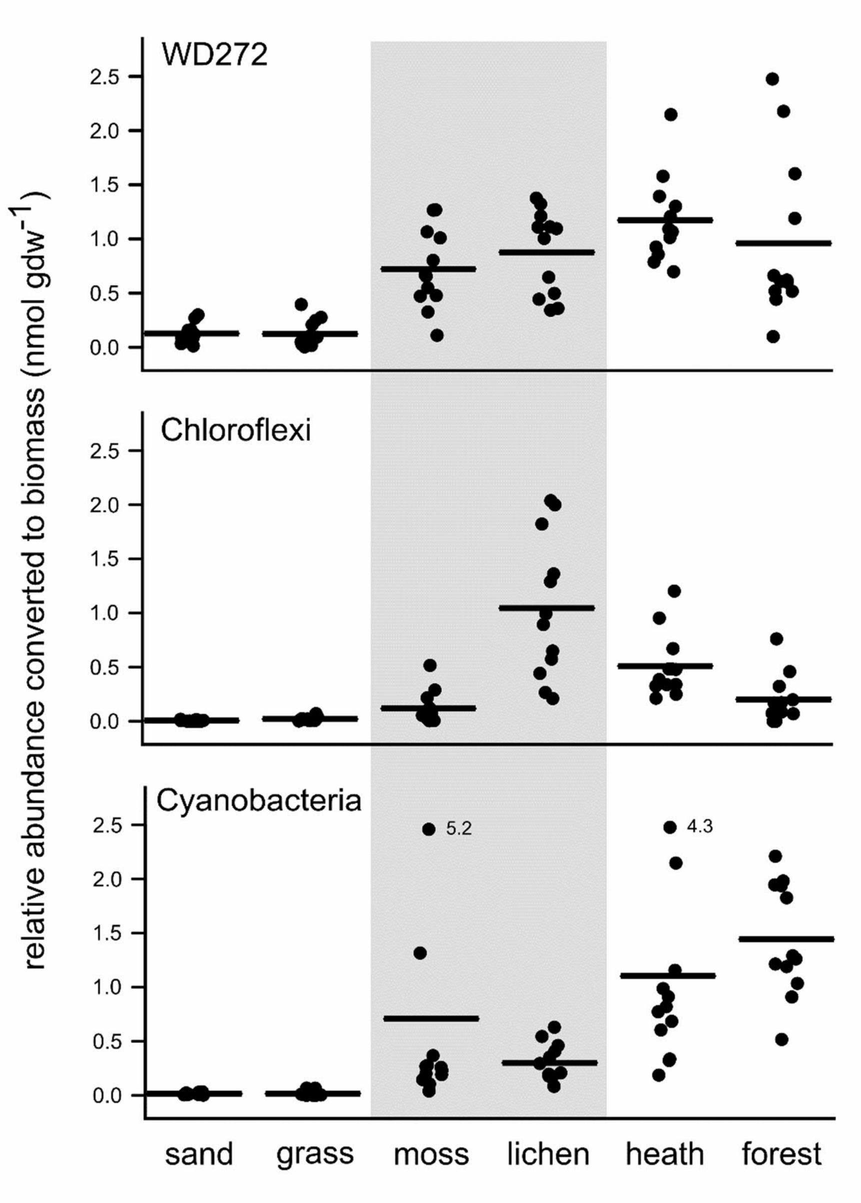


**Figure S5** Biomass estimates of selected early and cryptogam stage bacteria with sand dune successional stage. Relative abundances from sequencing data were converted into biomass based on total microbial biomass in PLFA analysis. Horizontal lines indicate mean values. Labelled points are outliers that would be outside the plot area. Note that primer bias and taxon-specific differences in DNA extraction efficiency can affect the conversion of sequencing results to biomass.

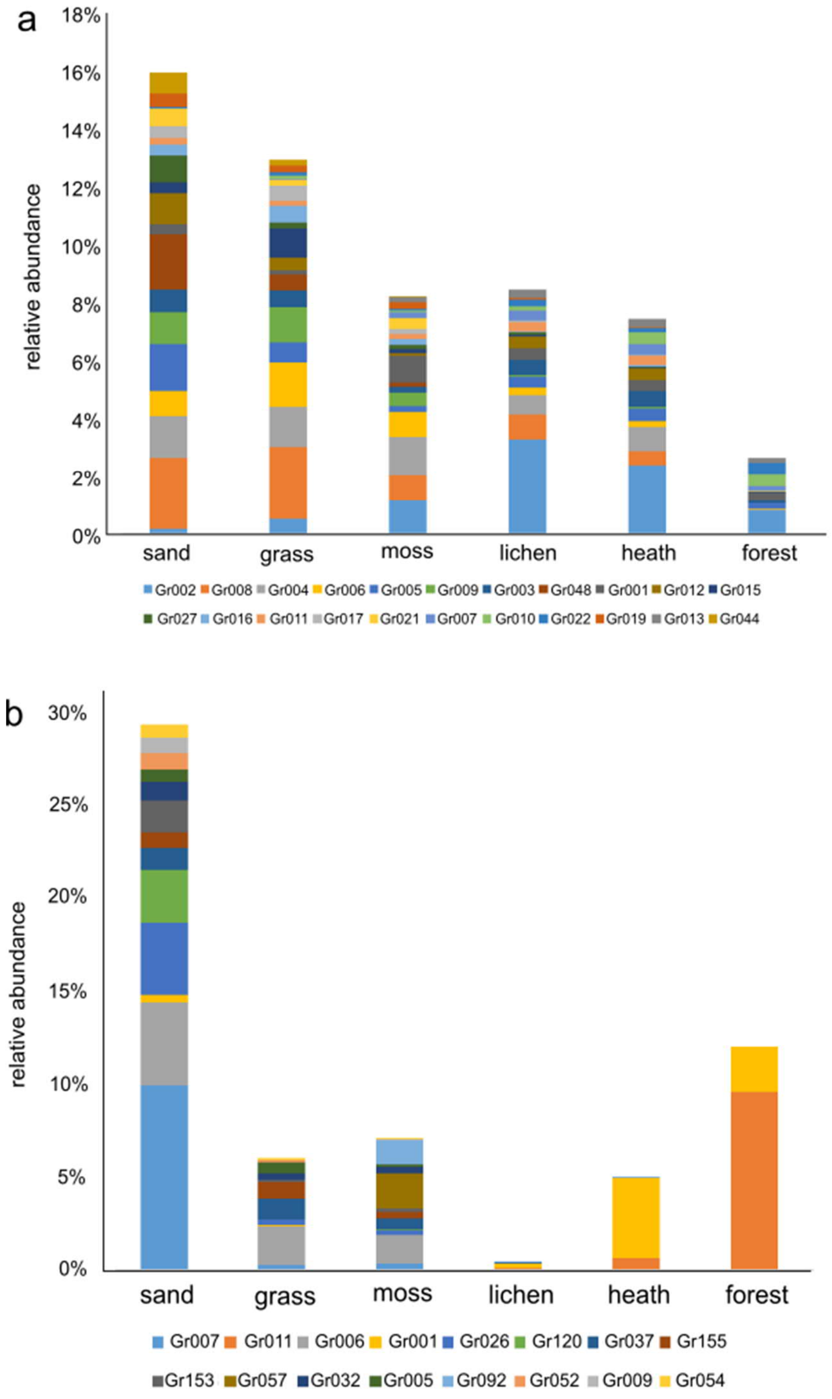


**Figure S6** Unclassified (A) bacterial phylum level groups (75% 16S rRNA gene sequence similarity) and (B) fungal class level groups (75% ITS2 sequence similarity) as percentage of total sequence reads per sand dune successional stage (four deflation basins and three replicate samples combined per stage). Groups with relative abundance of >0.15% of total reads are included.

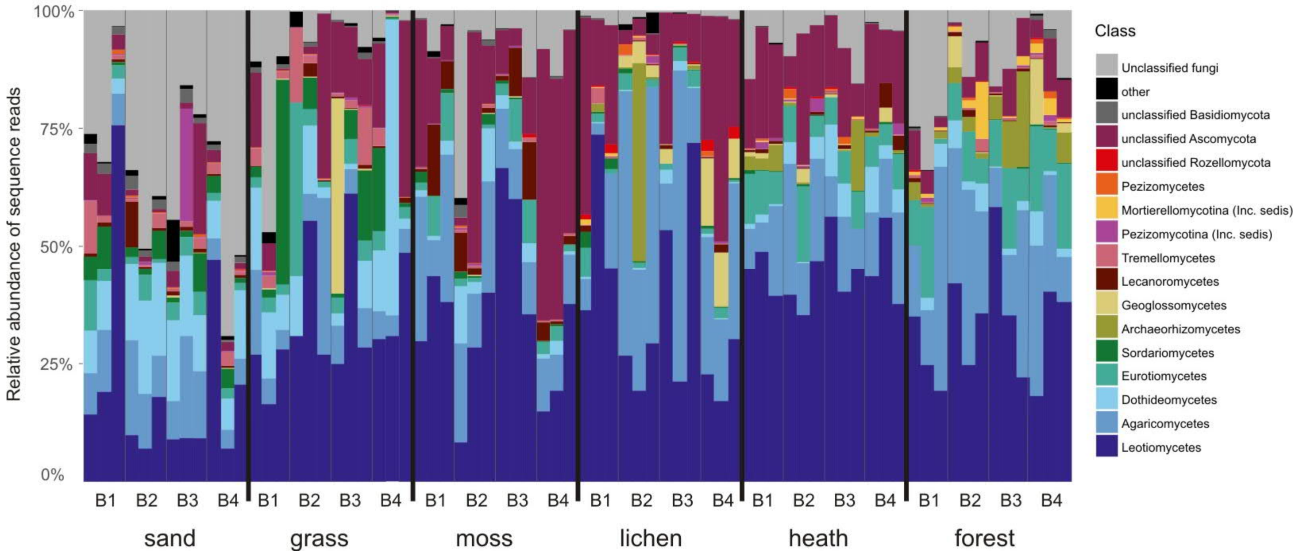


**Figure S7** Class-level community composition of fungi based on ITS2 region sequencing of six sand dune successional stages. B1-B4 indicate four deflation basins, and three columns within a deflation basin are replicate soil samples. ‘Other’ includes Microbotryomycetes, Mucoromycotina (Incertae sedis), Taphrinomycetes, Pucciniomycetes, Ustilaginomycotina (Incertae sedis), Chytridiomycetes, Exobasidiomycetes, Ascomycota Incertae sedis, Glomeromycetes, Cystobasidiomycetes, Saccharomycetes, Orbiliomycetes, Wallemiomycetes, and Ustilaginomycetes. Inc. sedis, Incertae sedis.
